# Secreted LysM proteins are required for niche competition and full virulence in *Pseudomonas savastanoi* during host plant infection

**DOI:** 10.1101/2025.04.11.648327

**Authors:** Hilario Domínguez-Cerván, Laura Barrientos-Moreno, Luis Díaz-Martínez, Jesús Murillo, Inmaculada Pérez-Dorado, Cayo Ramos, Luis Rodríguez-Moreno

## Abstract

Phytopathogenic bacteria secrete diverse virulence factors to manipulate host defenses and establish infection. Characterization of the type III secretion system (T3SS)- and HrpL-independent secretome (T3-IS) in *Pseudomonas savastanoi* pv. *savastanoi* (Psv), the causal agent of olive knot disease, identified five secreted LysM-containing proteins (LysM1–LysM5) associated with distinct physiological processes critical for infection. Functional predictions from network analyses suggest that LysM1, LysM2, and LysM4 may participate in type IV pilus-related functions, while LysM3 and LysM5 are likely to possess peptidoglycan hydrolase domains critical for cell division. Supporting these predictions, loss of LysM1 function resulted in impaired twitching and swimming motility, highlighting a role in pilus-mediated movement and early host colonization. In contrast, mutants lacking LysM3 or LysM5 exhibited pronounced filamentation and defective bacterial division, underscoring their essential role in septation, a process crucial for both *in planta* fitness and tumor formation. Structural modeling and protein stability assays demonstrate that LysM3 interacts with peptidoglycan fragments such as tetra-N-acetylglucosamine and meso-diaminopimelic acid, as well as with zinc ions, through conserved LysM and M23 domains. LysM3 also displayed selective bacteriostatic activity against co-inhabiting Gram-negative bacterial competitors, such as *Pantoea agglomerans* and *Erwinia toletana*. Our findings highlight the relevance of LysM proteins in maintaining bacterial integrity, motility, and competitive fitness, which are crucial for successful host infection. This study expands the functional repertoire of LysM-containing proteins and reveals their broader impact on bacterial virulence and adaptation to the plant-associated niche.

**Author Summary:** Plant pathogenic bacteria secrete a variety of proteins to manipulate host responses and outcompete microbial rivals. In this study, we investigated a group of five secreted proteins with conserved LysM domains produced by the bacterium responsible for olive knot disease, *Pseudomonas savastanoi*. We found that these proteins fulfill distinct and critical functions during infection. Three are potentially involved in the assembly of type IV pili, structures that help bacteria move and colonize plant tissues, while the other two are related to enzymes that remodel the bacterial cell wall and are essential for proper cell division. Mutants lacking these proteins failed to divide normally and were significantly impaired in their ability to infect olive plants. One of these proteins, LysM3, also exhibited the ability to inhibit other Gram-negative bacteria that coexist with the pathogen in the plant tumor, pointing to a role in microbial competition. These findings shed light on how LysM-containing proteins contribute to both disease progression and bacterial competition and survival in the plant environment. This work paves the way for new insights into bacterial pathogenesis and offers potential strategies for controlling olive knot disease and other plant infections.

## Introduction

The plant apoplast is the extracellular space encompassing the fibrillar matrix of cell walls, intercellular fluid-filled spaces, the xylem-lumen, the cuticle covering the outer plant surface, and air-filled regions (1). It plays a critical role in various physiological processes, including long-distance intercellular signaling, water and nutrient transport, and the perception of biotic stresses. As part of their preformed defense mechanisms, plants secrete hydrolytic enzymes such as chitinases and peptidases into the apoplast. These enzymes degrade conserved structural components of fungal and bacterial cell walls, thereby triggering immune responses (2–5). Plant cells monitor the apoplast through membrane-bound immune receptors that detect microbially-derived molecules or signs of microbial activity within plant tissues (6,7). Common examples of these molecules include fungal chitin (8) and bacterial peptidoglycan (9), both of which are recognized by plant immune receptors containing lysin motifs (LysM) (3,10–12).

LysMs are a class of carbohydrate-binding modules (CBMs) that specifically recognize and bind polysaccharides containing N-acetylglucosamine (GlcNAc) residues (13). These functional units are ubiquitous across all domains of life except Archaea, participating in very different cellular processes. Besides their occurrence in immune receptors, in particular, their role in pathogenicity has been extensively characterized in fungal pathogens (14–17). Fungal LysM effectors contribute to plant infection by preventing host recognition of chitin oligomers (18) or shielding fungal cell walls from host chitinases (19). It is not known, however, if these domains are also relevant virulence determinants for bacterial plant pathogens, where most identified LysM-containing proteins are peptidoglycan (PG) hydrolases, which are essential for cell-wall remodeling, bacterial growth and interaction with the host (13,20).

Phytopathogenic bacteria enter plant tissues through natural openings or wounds and establish infection using diverse strategies, including the deployment of specialized secretion systems that translocate virulence factors into the host (21–23). Among these, the type III secretion system (T3SS) is a major virulence determinant of Gram-negative phytopathogenic bacteria (24). The T3SS enables direct injection of effector proteins (T3Es) into the host cytoplasm, where they manipulate host cellular processes and suppress immune responses (25–27). Similarly, type IV (T4SS) and type VI (T6SS) secretion systems facilitate the translocation of other groups of bacterial effectors across both the inner and outer membranes into eukaryotic or bacterial target cells (28,29). Other secretion systems, such as type I (T1SS), type II (T2SS), and type V (T5SS), contribute to bacterial virulence by secreting proteins and metabolites into the extracellular space during host colonization. The secreted factors include cell-wall degrading enzymes, proteases, lipases, adhesins, phosphatases and carbohydrate-processing proteins (30–32). However, although large amounts of putatively secreted factors have been identified through genome annotations, it is currently unknown which factors are actually secreted and play roles in the interaction with plant hosts during infection (32). Characterization of the secretomes of phytopathogenic bacteria is thus essential for understanding the molecular mechanisms underpinning niche establishment and disease progression.

The *Pseudomonas syringae* complex is one of the most significant groups of phytopathogenic bacteria due to their diseases caused in agricultural and ornamental crops. Among them, strains of *Pseudomonas savastanoi* pv. savastanoi (Psv), the causal agent of olive knot disease (33), induce hyperplastic growth in host tissue through the coordinated action of multiple virulence factors (34–36). A key determinant of Psv virulence is the T3SS, which translocates effectors into host cells. In Psv NCPPB 3335, deletion of *hrpA*, encoding the structural subunit of the T3SS pilus, impairs tumor formation and bacterial colonization of host tissues (37). Similarly, deletion of *hrpL*, a key regulator of T3SS effectors, structural components and other virulence-related genes, disrupts effector secretion and attenuates virulence (38). However, *hrpA* and *hrpL* mutants remain viable within host tissue, highlighting the potential role of proteins secreted independently of the T3SS and HrpL that act in bacterial virulence and fitness.

In this work, we analyzed the T3SS- and HrpL-independent secretome of Psv and identified five secreted LysM domain-containing proteins, which we named LysM1 to LysM5. Three of these (LysM1, LysM2 and LysM4) are putatively involved in type IV pilus (T4P) assembly, whereas LysM3 and LysM5 are putative PG hydrolases crucial for bacterial fitness and physiology. Loss-of-function mutations in these genes impaired bacterial colonization of plant tissues and attenuated bacterial virulence. Specifically, *lysM3* and *lysM5* mutations caused severe defects in cell septation, compromising bacterial fitness both *in vitro* as well as during plant infection. Structural modelling and Nano Differential Scanning Fluorimetry (nanoDSF) assays showed that LysM3 harbors an active LysM domain and an M23 peptidase domain, while bacterial growth assays demonstrated its selective bacteriostatic activity against Gram-negative bacteria, including *Pantoea agglomerans* and *Erwinia toletana*, which co-inhabit olive tumors with Psv. These findings underscore novel roles of LysM proteins in bacterial fitness and virulence, as well as in competition with the bacterial communities in diseased plant tissues, emphasizing their broader relevance to the pathogenicity of bacterial phytopathogens.

## Results

### Identification of the T3SS-independent secretome in *Pseudomonas savastanoi* pv. savastanoi NCPPB 3335

We determined the secretome of Psv NCPPB 3335 in HIM medium (39), a minimal medium that mimics conditions in the plant apoplast and induces the expression of bacterial virulence genes (39). The complex proteome of Psv was analyzed using mass spectrometry (MS) in combination with bioinformatics. MS outputs at 12 and 24 hours post-incubation (hpi) were processed through our custom-designed bioinformatics pipeline, SecretFlow, to identify extracellular predicted proteins (EPPs) for each sample.

To identify proteins that are secreted independently of the T3SS and HrpL, EPPs from the wild-type Psv strain were compared with those from *hrpA* and *hrpL* mutants (Psv-ΔhrpA, and Psv-ΔhrpL, respectively) at 12 and 24 hpi (Figure 1). At 12 hpi, a total of 429, 163 and 163 EPPs were identified for Psv NCPPB 3335, Psv-ΔhrpA, and Psv-ΔhrpL, respectively (Figure 1A). Similarly, at 24 hpi, 771, 637 and 327 EPPs were identified for the respective Psv strains (Figure 1B). Proteins common to the secretomes of all three strains were considered part of the core T3SS-independent and HrpL-independent secretome (T3-IS).

**Figure 1.**
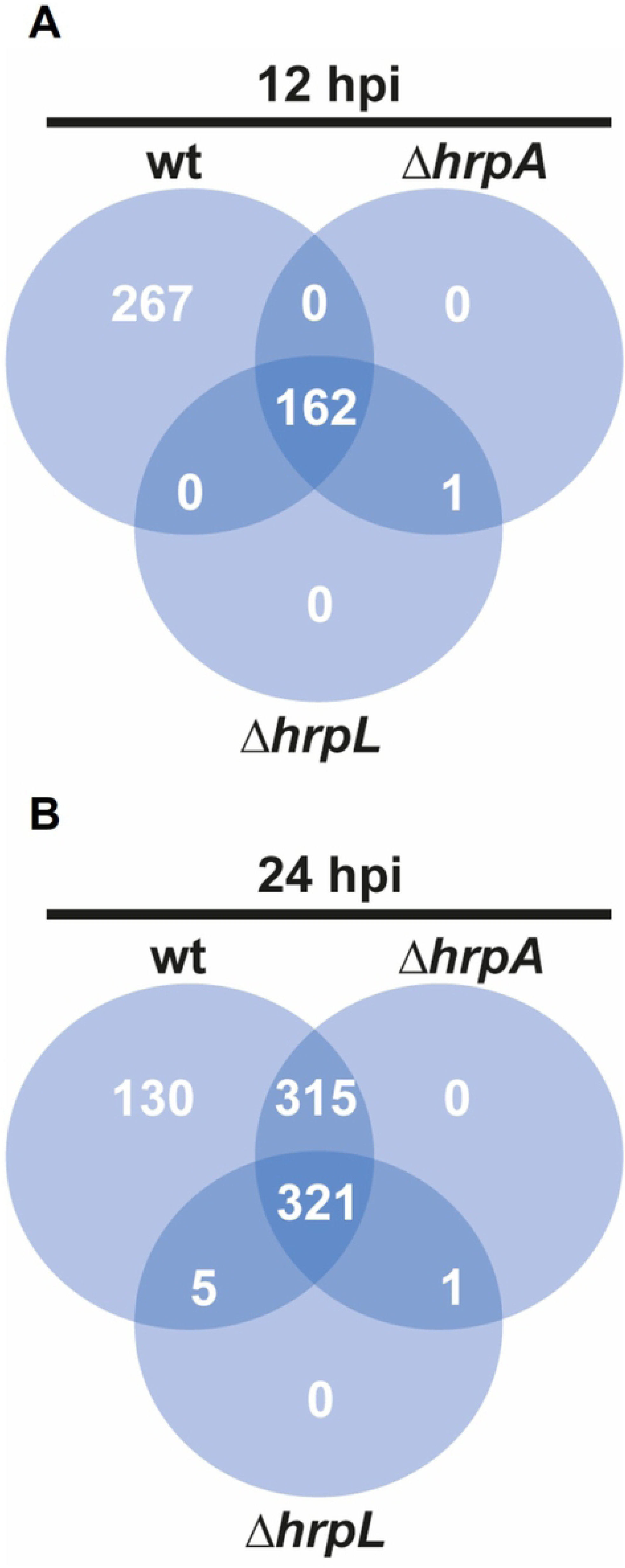
Extracellular proteins in apoplast-mimicking minimal (HIM) medium. Venn diagrams show the extracellular proteins identified in the secretome of the wild-type strain *P. savastanoi* pv. savastanoi NCPPB 3335 (wt), and its Δ*hrpA* and Δ*hrpL* mutants, lacking structural and regulatory components of the type III secretion system, respectively, after 12 hours (A) and 24 hours (B) of incubation in HIM medium. At 24 h, 96.3% of the proteins shared by all three strains were also detected at 12 h.

A total of 162 and 321 proteins comprised the T3-IS at 12 and 24 hpi, respectively (Figure 1A and 1B). Most of the proteins identified at 12 hpi (156 proteins, 96.3%) were also found among the 321 proteins identified at 24 hpi. Detailed information on these 321 EPPs is provided in Supporting Information Table S1. Given the higher number of EPPs identified at 24 hpi, subsequent analyses focused on the T3-IS from this time point.

### Functional categorization and in-depth analysis of the T3SS-independent secretome

The T3-IS was functionally categorized using the ShinyGO software (40), with annotations from the GO molecular functions, GO biological processes, and Uniprot databases. This analysis classified the T3-IS into 48 distinct molecular categories (Figure 2). The combined use of these three databases covered 57% (183 proteins) of the T3-IS from Psv. Among the most represented categories were periplasmic space, envelope, transporter activity, response to stimulus, macromolecule localization, peptidase activity, molecular transducer activity, signaling receptor activity, channel activity, and carbohydrate binding (Figure 2). We performed a functional enrichment analysis using ShinyGO, comparing the abundance of GO terms in the secretome compared to their abundance in the complete deduced proteome of Psv NCPPB 3335 (Figure 2). The functional enrichment analysis revealed differences between 12 hpi and 24 hpi (Figure S1 and Figure 2), although the most enriched categories at both time points were peptide and amino acid binding, lipid transport, and specific enzymatic activities, such as aldose-1 epimerase and serine-type carboxypeptidase or exopeptidase activities.

**Figure 2.**
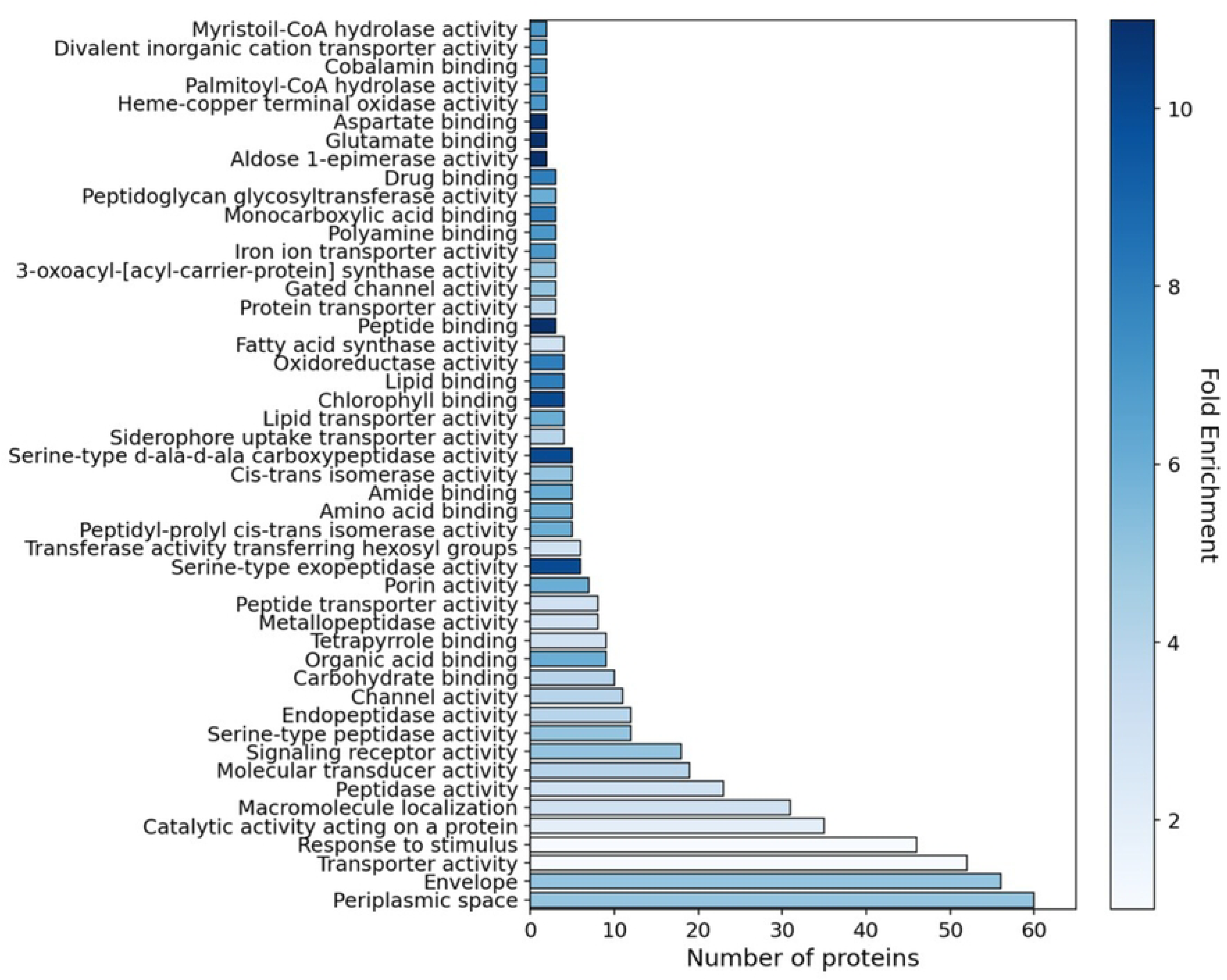
Functional enrichment analysis of the *Pseudomonas savastanoi* NCPPB 3335 secretome using ShinyGO. The secretome was obtained after 24 hours of incubation in HIM medium. The analysis highlights significantly enriched GO functional categories, classified by molecular function and cellular localization. The X-axis represents the number of proteins associated with each category, while the color indicates the degree of enrichment (fold enrichment). Fold enrichment reflects the frequency of a given GO term in the secretome compared to its frequency in the complete deduced proteome of strain NCPPB 3335. The analysis was performed with a false discovery rate (FDR) cutoff of 0.05.

Given the role of carbohydrate recognition in growth, cell adhesion and signaling in living organisms, we focused on proteins containing conserved carbohydrate-binding modules (CBMs). Within this group, we identified five proteins featuring LysM domains (LysM1 to LysM5; Figure 3A). Except for LysM5, which has its LysM domain located at the C-terminus, the other four proteins exhibited a single LysM domain at the N-terminus. InterProScan analysis revealed additional functional domains: LysM1 contains a tetratricopeptide domain, which facilitates protein-protein interactions, playing key roles in processes such as cell cycle regulation, transcription, and protein transport (41); LysM1 and LysM2 share a FimV domain at their C-terminus, involved in the biogenesis and function of bacterial pili, critical for adhesion, motility, and host colonization (42). LysM3 features an M23-type peptidase domain at its C-terminus, found in diverse zinc metallopeptidases that hydrolyze the bacterial peptidoglycan, contributing to cell wall remodeling and virulence (43). LysM5 includes an amidase N-terminal (AMIN) domain, found in amidases that degrade bacterial cell walls, promoting cell division and lysis (44), and an N-acetylmuramoyl-L-alanine amidase domain, which catalyzes peptidoglycan hydrolysis, aiding bacterial cell wall remodeling during cell division and host immune evasion (45) (Figure 3A).

**Figure 3.**
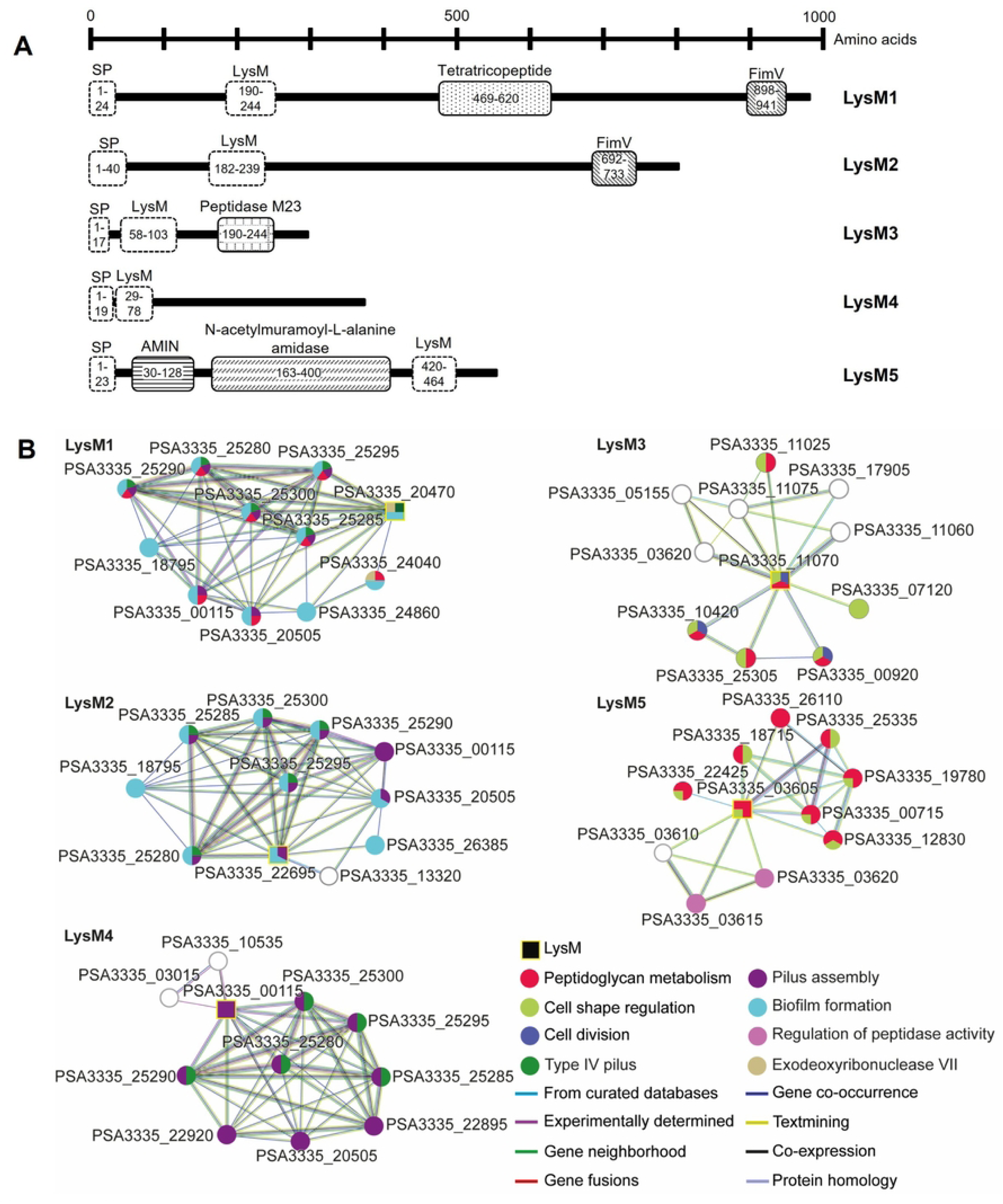
Characteristics of LysM proteins. (A) Schematic representation of LysM proteins and their functional domains identified using InterProScan. The diagram displays the structural organization of the LysM proteins, including predicted domains and regions of interest, with numbers indicating their position in the sequence (LysM1 Refseq ID: WP_031594872.1; LysM2: WP_002552571.1; LysM3: WP_002554701.1; LysM4: WP_002551334.1; LysM5: WP_199523490.1). Dashed lines indicate conserved domains or features present in all proteins, while filled patterns represent domains that are either exclusive to a single protein or shared among only some of them. Notably, SP represents the signal peptide while AMIN corresponds to an N-terminal amidase domain. (B) STRING interaction network analysis showing the functional relationships between LysM proteins and the products of other genes, providing insights into the potential functions of LysM proteins. The color of the nodes represents the functional category to which each protein belongs, while the color of the edges indicates the source of the interaction.

To predict the potential functions of the five LysM proteins, we conducted a predictive relationship analysis using the STRING software (Figure 3B). This analysis suggested that the LysM proteins can be divided into two distinct subgroups: LysM1, LysM2, and LysM4, which are likely involved in type IV pilus assembly, biofilm formation, and cell membrane modification (Figure 3B), and LysM3 and LysM5, which are predicted to play roles in cell division and the maintenance of cell shape (Figure 3B).

### Influence of LysM1, LysM2, and LysM4 proteins in motility

The potential involvement of LysM1, LysM2, and LysM4 in type IV pilus assembly (Figure 3B) along with the FimV domains found in the LysM1 and LysM2 sequences, which are associated with type IV pilus (Figure 3A), prompted us to investigate their role in bacterial motility. Twitching motility is a type IV pili-dependent but flagella-independent movement, whereas flagella are responsible for swimming motility (46). Given the involvement of both motility types in adhesion and host colonization, twitching and swimming motility were assessed in the Δ*lysM1*, Δ*lysM2*, and Δ*lysM4* mutants (Figure 4).

**Figure 4.**
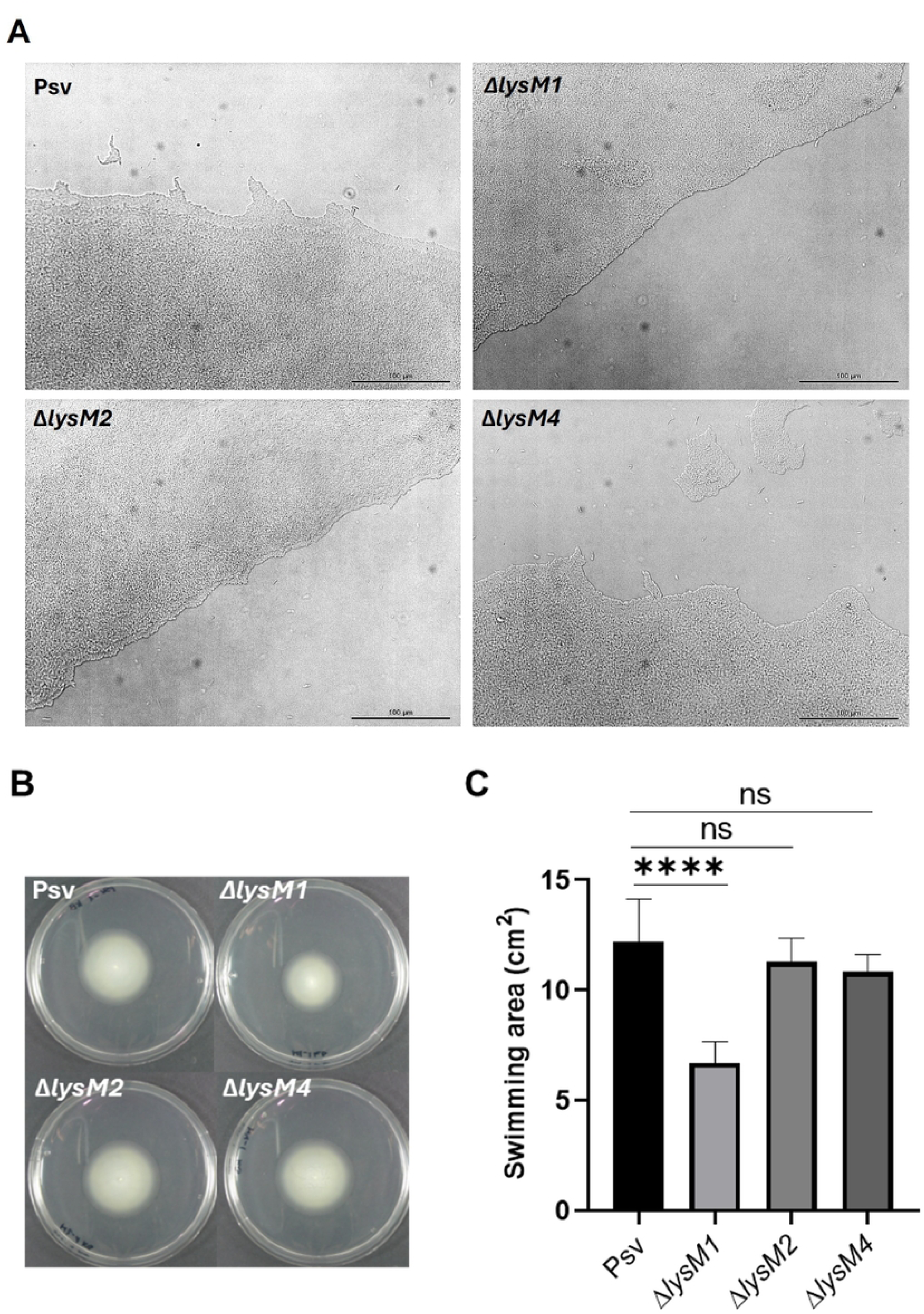
Impact of Δ*lysM* mutations on bacterial motility. (A) Twitching motility on 1%-agar KB medium overlaid on slides by the wild-type strain Psv NCPPB 3335 (Psv) and the Δ*lysM* mutants. Microscopy images were taken after 24h of incubation at 25°C with Nikon Eclipse E800 light microscope (20x). The experiment was repeated three times with two repetitions per trial, and a representative image is shown per strain. (B-C) Swimming motility on 0.3%-agar KB medium. (B) Representative images of swimming halos formed by the wild-type strain Psv and the Δ*lysM* mutants after three days of incubation at 25°C. (C) Swimming halos are represented as the covered area (cm^2^). Data are presented as means and standard deviation (SD) from three independent experiments, with at least four replicas per assay. Asterisks indicate statistically significant differences (ANOVA test; p < 0.0001), and “ns” indicates no significant differences between samples.

Twitching motility was evaluated on a solid surface under humid conditions, and twitching areas were visualized with light microscopy. As shown in Figure 4A, the wild-type strain exhibited the characteristic irregular or poorly defined edge at the leading front of the movement, with well-organized cell clusters and single cells at the forefront of motility. Similarly, the Δ*lysM2* and Δ*lysM4* mutants behaved as the wild-type strain. Notably, the defective Δ*lysM2* strain formed smaller and less pronounced protrusions than the wild type. In contrast, the strain containing mutation Δ*lysM1* displayed an almost flat and well-defined advancing front, suggesting alterations in twitching motility dynamics. These findings indicate that LysM1 may play a role in the functionality of the type IV pili.

Swimming motility was assessed under low-viscosity conditions [0.3% agar (wt/vol)] in KB medium. The swimming halos produced by the Δ*lysM1* mutant were significantly smaller than those of the wild-type strain (Figure 4B, C). However, the strains with the Δ*lysM2* and Δ*lysM4* mutations did not show any defects in swimming motility under these conditions.

### LysM3 and LysM5 are involved in bacterial cell septation during cell division in Psv NCPPB 3335

LysM domains are involved in various aspects of the bacterial life cycle (13). Additionally, the peptidase M23 and N-acetylmuramoyl-L-alanine amidase domains found in LysM3 and LysM5, respectively, are potentially involved in peptidoglycan hydrolysis to facilitate cell wall expansion and organization during bacterial growth (45,47). Therefore, we compared the growth profiles of the LysM mutants to that of the wild-type strain.

No significant differences were observed in growth patterns or maximal optical density at 600 nm (OD_600_) values between the wild-type strain and the *lysM* mutants in rich KB medium (Supplementary Figure 2A). However, the Δ*lysM5* mutant consistently exhibited a slight, albeit no significant, trend toward lower OD_600_ values compared to the wild-type strain and the other mutants. In contrast, after 24 hours of growth in KB, the colony-forming unit (CFU) counts for the defective Δ*lysM3* and Δ*lysM5* strains were significantly lower than those of the wild-type strain and the remaining mutants (Supplementary Figure 2B).

Given the reduced CFU counts for the strains containing mutations Δ*lysM3* and Δ*lysM5* (Supplementary Figure 2B) and the functional predictions from the STRING analysis obtained for these proteins (Figure 3B), we further investigated cell morphology of all the mutants and their complemented strains using confocal microscopy. Notably, the defective Δ*lysM3* and Δ*lysM5* strains exhibited a distinctive phenotype, characterized by unsegmented chains of cells joined at their polar ends (Figure 5A), likely due to impaired septum formation during cell division. In contrast, the Δ*lysM1*, Δ*lysM2*, and Δ*lysM4* knockout strains displayed typical rod-shaped morphology similar to the wild-type strain. The filamentous phenotype was rescued in the complemented Δ*lysM3* and Δ*lysM5* mutant strains, which displayed normal cell morphology (Figure 5A).

**Figure 5.**
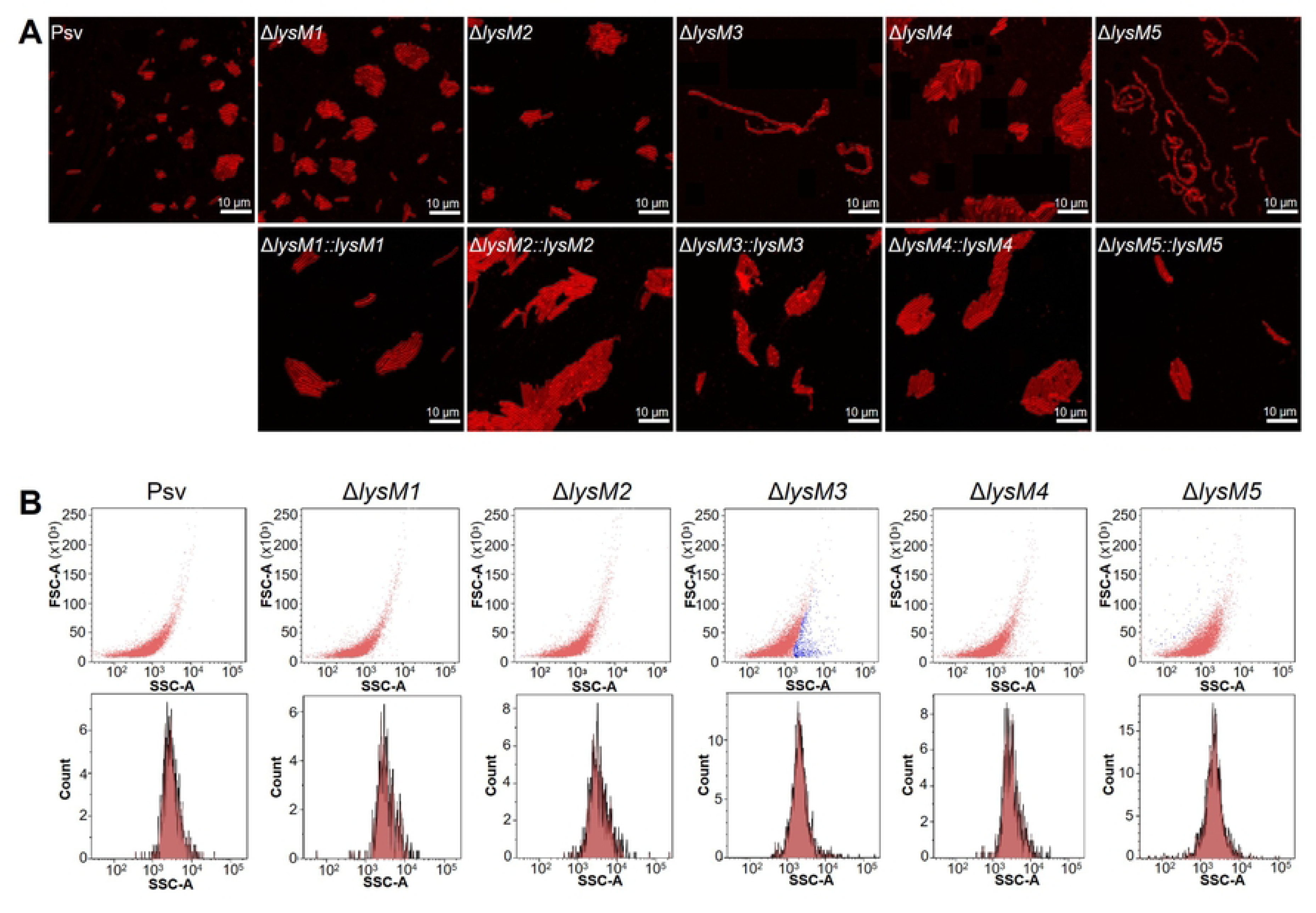
Lack of LysM3 and LysM5 causes altered cell morphology in *P. savastanoi* pv. savastanoi NCPPB 3335. (A) Confocal microscopy images of the wild-type strain and each of the five Δ*lysM* mutants, along with their corresponding complemented strains. Cells were stained with the membrane-binding fluorescence dye FM4-64 (red). (B) Scatter plots (upper plots) illustrate the relationship between cell size (FSC-A, forward scatter) and cell complexity (SSC-A, side scatter). Each point represents an individual event (cell), with the X- and Y-axes showing complexity and size, respectively. The blue-colored dots in the scatter plots represent subpopulations that exhibit differences in size or complexity compared to the rest of the bacterial population. Histograms (lower plots) show the distribution of the number of events (Count, as log_10_ of number of cells) as a function of cell complexity (SSC-A). The X-axis represents cell complexity, while the Y-axis shows the represents of events.

Flow cytometry analysis of size and complexity in exponentially growing cultures supported the phenotypes observed by confocal microscopy (Figure 5B). Comparative analysis revealed that a subset of the Δ*lysM3* knockout strain population exhibited greater complexity compared to the other mutants and the wild-type strain. Similarly, the defective Δ*lysM5* strain showed a subpopulation with increased cell size, which was absent in the other strains (Figure 5B). Furthermore, the number of complex events was significantly higher in the Δ*lysM3* and Δ*lysM5* mutants compared to the wild-type and the other mutants, consistent with their filamentous morphology.

### Extracellular LysM proteins contribute to *P. savastanoi* fitness and virulence in olive plants

Given the well-established role of LysM domains in fungal virulence (15,48), we evaluated the virulence of mutants lacking each of the five secreted LysM-containing proteins. *In vitro* grown olive plants provided a rapid and sensitive model to assess olive knot disease symptoms, which fully develop within 30 days post-inoculation (dpi) (Figure 6A). At 30 dpi, the Δ*lysM1*, Δ*lysM2*, Δ*lysM4*, and Δ*lysM5* mutants induced tumors at the inoculation sites that were morphologically similar to those produced by the wild-type strain. However, tumor volumes were significantly reduced for all mutants compared to the wild-type strain, with more pronounced reductions observed for tumors induced by the defective Δ*lysM1* and Δ*lysM3* strains (Figure 6B). Notably, plants inoculated with the strain containing mutation Δ*lysM3* developed only small outgrowths that failed to progress into mature tumors (Figure 6A).

**Figure 6.**
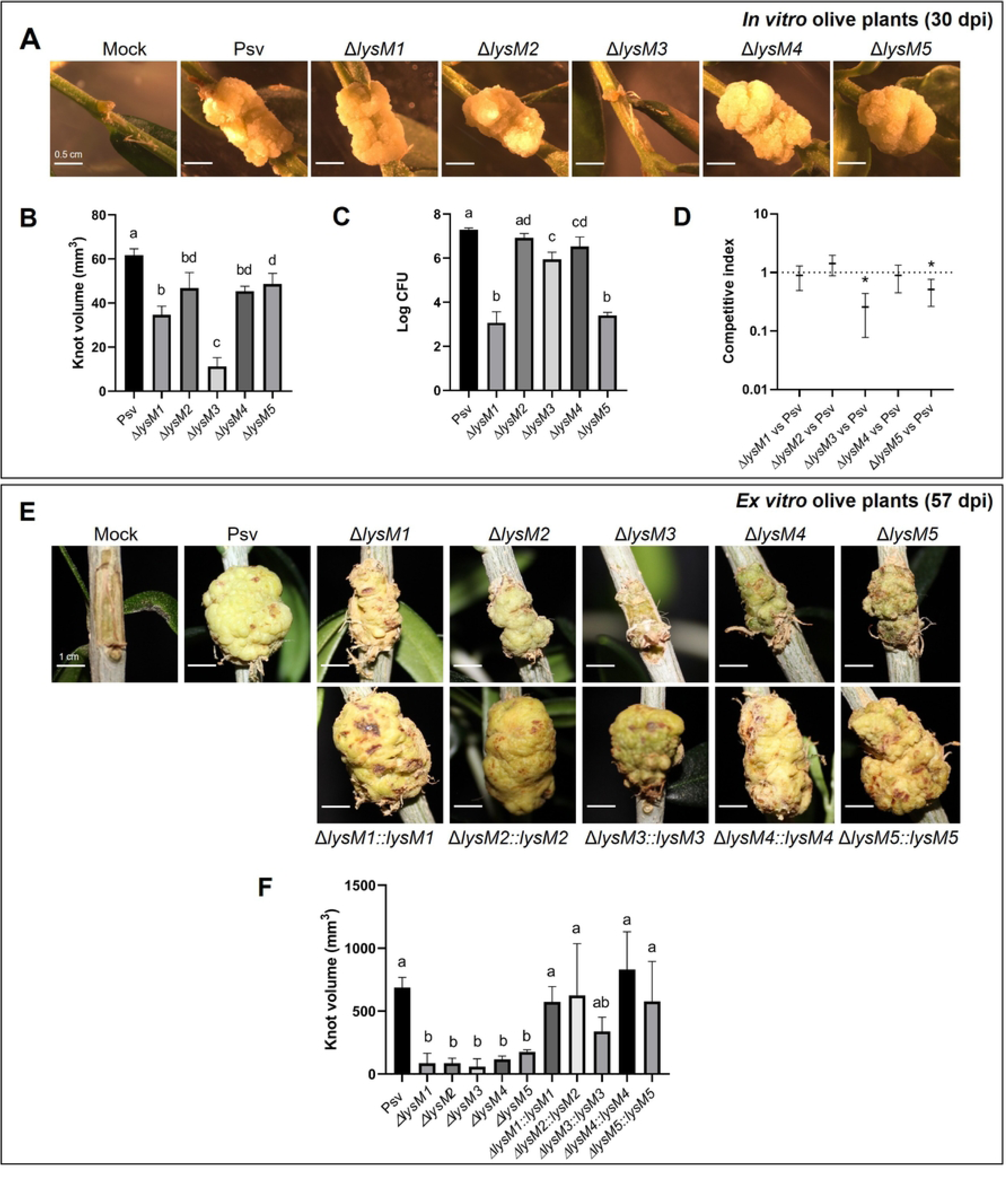
Virulence and competitive fitness of *P. savastanoi lysM* mutants in olive plants. (A) Representative symptoms induced by the wild-type strain *P. savastanoi* pv. savastanoi NCPPB 3335 and its five *lysM* mutants. Mock, plant inoculated with buffer. (B) Total bacterial populations recovered from inoculated tissues. (C) Quantification of knot volume induced by the indicated strains. (D) Competitive index for mixed 1:1 inoculations. An asterisk denotes a value significantly different from one, as determined by a Student’s t-test (p = 0.05). (E) Representative symptoms induced by the wild-type strain, the Δ*lysM* mutants and the complemented strains. Mock, plant inoculated with buffer. (F) Quantification of knot volume induced by the indicated strains. Data for panels A-D were collected 30 days post-inoculation of *in vitro* olive plants, while data for panels E-F were collected 57 days post-inoculation of *ex vitro* olive plants. Bars and data points (panel D) represent means from five biological replicates (four for panel D) with standard error. Different letters indicate statistically significant differences, as determined by ANOVA followed by Tukey’s t-test (p < 0.05). Scale bars = 0.5 cm (A) and 1 cm (E). Complemented strains are indicated by the symbol ‘::’ in the strain names, followed by the corresponding gene.

To evaluate bacterial colonization, we quantified CFU counts in infected plants at the end of the experiment, which broadly aligned with disease severity but did not show a strict correlation. Specifically, while the defective Δ*lysM2*, Δ*lysM3*, and Δ*lysM4* strains showed slight but significant reductions in CFU counts compared to the wild-type strain, the Δ*lysM1* and Δ*lysM5* mutants exhibited severely impaired colonization of *in vitro* olive tissues (Figure 6C). To determine whether the reduced colonization ability of the mutants also resulted in decreased competitive fitness, we performed competition assays against the wild-type strain. While the Δ*lysM1*, Δ*lysM2*, and Δ*lysM4* knockout mutants showed no significant differences in competitive fitness, the strains containing mutations Δ*lysM3* and Δ*lysM5* were significantly outcompeted by the wild-type strain (Figure 6D).

Since *in vitro* olive plants lack autotrophy and lignified tissues, which may facilitate symptoms development, we also conducted virulence assays using *ex vitro* olive plants with all the Δ*lysM* mutants and their complemented strains (Figure 6E). In this system, all Δ*lysM* knockout strains exhibited significantly reduced tumor volumes compared to the wild-type strain after 57 dpi (Figure 6E and Figure 6F). Complementation of each mutant strain restored tumor appearance (Figure 6E) and size (Figure 6F) to levels comparable to those of tumors induced by the wild-type strain, demonstrating that the virulence defects observed for the mutants were specifically due to the *lysM* gene deletions.

### LysM3 binds bacterial peptidoglycan fragments and analogues via conserved functional domains

Given its critical role in symptoms development (Figure 6), we further characterize LysM3, a putative PG DD-metalloendopeptidase containing a LysM domain and a peptidase M23 domain (Figure 3A), suggesting its interaction with the peptidoglycan. To investigate this, we compared the predicted structure of LysM3 with crystallized proteins containing LysM or M23 domains and conducted nanoDSF assays with purified LysM3 to evaluate potential ligand interactions (Figure 7).

**Figure 7.**
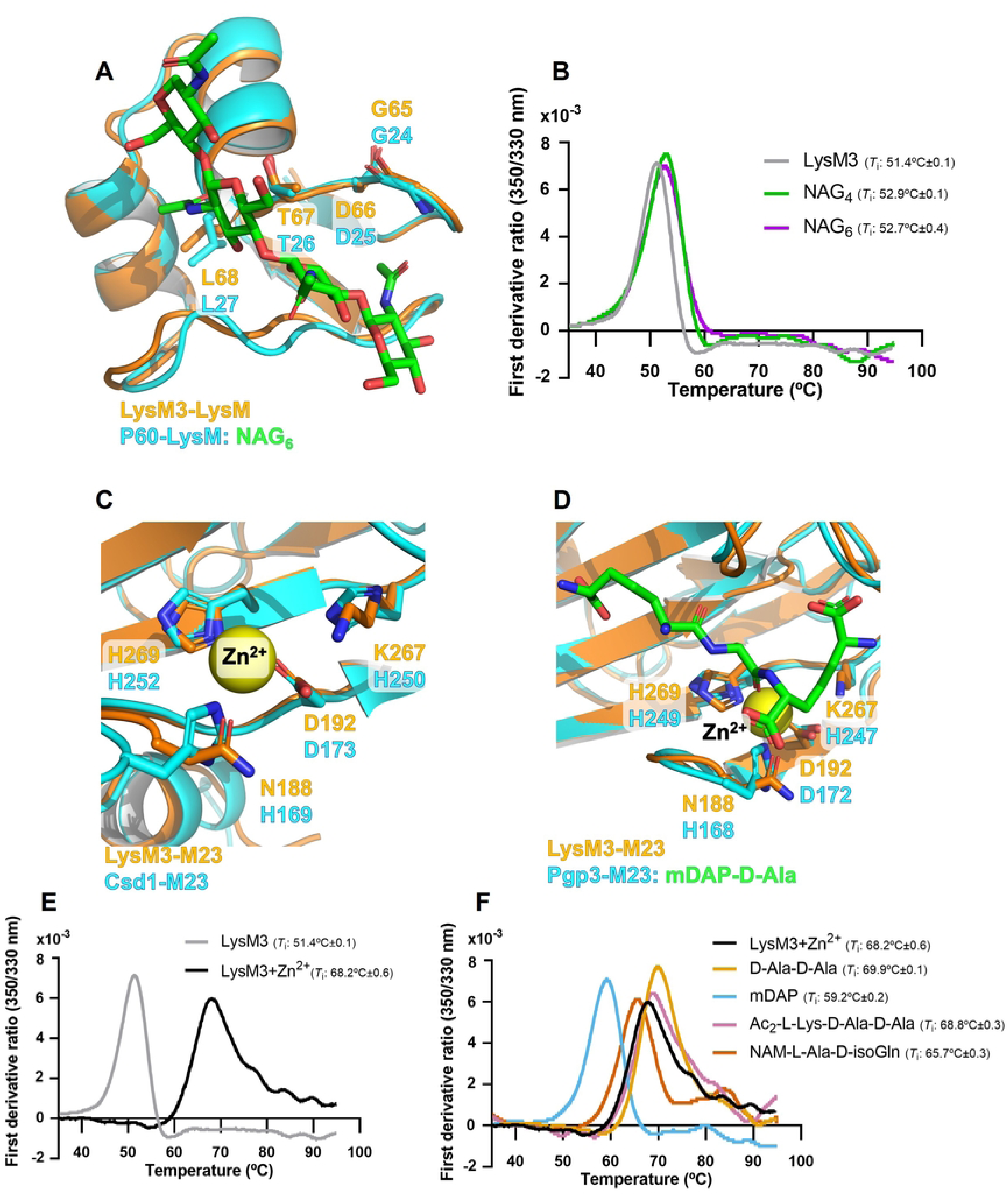
Interaction of LysM3 with peptidoglycan (PG) fragments/analogues. (A) Structural superimposition of LysM3-LysM domain (AlphaFold model, in orange) with the crystallographic complex of the LysM domain-containing protein P60 bound to NAG₆ (in cyan). The image highlights conservation of PG-binding features in LysM3-LysM domain. (B) Nano differential scanning fluorimetry (nanoDSF) analysis of LysM3 without Zn^2+^ in the presence of NAG₄ and NAG₆, showing its interaction with peptidoglycan analogues. Mean *T*_i_ values ± standard deviation (SD) is shown. (C-D) Superimposition of the M23 peptidase domain of LysM3 (Alphafold model, in orange) with the peptidase domains of Csd1 *apo* (in cyan) and Pgp3 bound to mDAP-D-Ala PG fragments (in cyan), showing conservation of residues involved in the coordination of the catalytic Zn^2+^ and in substrate interaction. (E) NanoDSF analysis of LysM3 showing its stabilization in the presence of Zn^2+^ compared to the protein alone. (F) Thermal stability profiles of LysM3 with Zn^2+^ and four PG fragments showing shifts of the *T*_i_ values in the presence of ligands. Mean *T*_i_ values ± SD are shown.

Comparison of the Alphafold-predicted LysM3 model (UniProt code: A0A0Q0APC2) with the experimental structure of functionally related proteins revealed conserved structural features. On the one hand, the LysM domain of LysM3 (LysM3-LysM) was superimposed with the P60 LysM domain solved in complex with hexa-N-acetyl-glucosamine (NAG_6_) (Protein Data Bank (PDB) code: 4uz3) (49). On the other hand, the M23 domain of LysM3 (LysM3-M23) was superimposed with the crystal structure of wild-type Csd1protein bound to zinc (PDB code: 5J1L) (50), and the Pgp3^H247A^ catalytically-inactive mutant protein, bound to meso-diaminopimelic acid-D-alanine (mDAP-D-Ala) (PDB code: 6jn1) (51).

LysM domains are characterized by βααβ fold and the presence of a highly conserved GDTL motif, commonly associated with the interaction of carbohydrate moieties of both fungal and bacterial cell walls (13). Superimposition of LysM3 and P60 models showed structural conservation of the βααβ fold and the GDTL motif, evidencing how LysM3 residues G65, D66, T67, and L68 of LysM3 overlay well with residues G24, D25, T26, and L27 of P60 (Figure 7A and Supplementary Figures 3A and 3B). Given this structural conservation, we conducted nanoDSF assays to examine interactions with peptidoglycan-derived oligosaccharides. NanoDSF measures protein stability by detecting changes in the inflection temperature (*T*_i_), associated to protein unfolding, where increases or decreases in *T*_i_ values indicate ligand interaction. Since peptidoglycan consists of alternating N-acetylmuramic (NAM) acid and N-acetylglucosamine units, we used NAG_4_ and NAG_6_ as ligand analogues for the nanoDSF experiments. Comparison of *T*_i_ values measured for LysM3 alone (51.4°C±0.1) and in the presence of NAG_4_ (52.9°C±0.1) and NAG_6_ (52.7°C±0.4) show an increase of >1°C in the presence of both oligosaccharides, which indicates interaction with both PG analogues (Figure 7B).

The M23 domain is present in zinc-dependent metallopeptidases, containing conserved HxxxD and HxH motifs providing a catalytic histidine and involved in the stabilization of the catalytic zinc (43). Structural superimposition of LysM3-M23 with Csd1 and Pgp3^H247A^ confirmed conservation of the general structure and key amino-acid residues involved in ligand binding (Supplementary Figure 3). Specifically, LysM3 residues D188 and H269 correspond to D192 and H252 (Supplementary Figure 3C), while variations at N188 and K267 in LysM3, corresponding to H169 and H250 in Csd1 (Supplementary Figure 3D and Figure 7C), indicated partial conservation of the active site. A similar scenario was observed with Pgp3^H247A^, where LysM3 residues D192 and H269 superimpose with D172 and H249 of Pgp3^H247A^ (Supplementary Figure 3E), while LysM3 variations at N188 and K267, corresponding to H168 and H247 in Pgp3 (Supplementary Figure 3F and Figure 7D), further indicated partial motif conservation.

Given the partial conservation of the HxxxD and HxH motifs in LysM3 and the known key role of Zn^2+^ in catalysis by M23 domains, we assess the interaction of LysM3 with Zn^2+^ using nanoDSF. The addition of Zn^2+^ resulted in a striking ∼20°C increase in the *T*_i_ (LysM3: 51.4°C±0.1; LysM3+Zn^2+^: 68.2°C±0.6), suggesting the interaction of Zn^2+^ at the active site (Figure 7E). To further explore the functionality of LysM3-M23, we tested the interaction of Zn^2+^-reconstituted LysM3 with peptidoglycan-derived ligands. All tested ligands caused variations in the *T*_i_ values (Figure 7F). Diacetyl-L-Lysine-D-Alanine-D-Alanine caused a modest increase in *T*_i_ value of 0.9°C, while D-Ala-D-Ala increased the *T*_i_ by 2°C. Conversely, N-acetylmuramyl-L-Alanine-D-isoglutamine decreased the *T*_i_ by 2°C, and meso-diaminopimelic acid (mDAP) caused a notable 9°C reduction in the *T*_i_, suggesting a strong interaction with LysM3-M23.

These results show that LysM3 retains functional features of LysM and M23 domains, interacting with both glycan and pentapeptide moieties of the PG. While the LysM domain exhibits classical peptidoglycan-binding properties, the M23 domain likely relies on Zn^2+^ coordination for activity.

### LysM3 exhibits selective bacteriostatic activity against Gram-negative bacteria

Proteins of the M23 family are involved in various processes, including cell division, maintenance of cellular structure, virulence, and, in certain organisms, lysing of cell walls of other bacteria for defense or feeding (47,52). Based on our nanoDSF analysis results, we evaluated the bacteriostatic activity of LysM3 against bacterial strains with diverse cell wall structures. We tested three Gram-negative bacterial strains: *E. coli* DH5α and two strains isolated from olive knots, *Erwinia toletana* DAPP-PG 735 (53) and *Pantoea agglomerans* DAPP-PG 734 (54). *Bacillus subtilis* 3610, a representative Gram-positive bacterium, was also included, with Psv NCPPB 3335, also isolated from olive knots, serving as the control strain.

Each strain was grown in rich medium alone (growth control) or supplemented with: buffer only (buffer control), buffer with a non-inhibitory protein (protein control), buffer with LysM4 (a protein containing only a LysM domain), buffer with LysM3 (containing both a LysM domain and a peptidase M23 domain), and buffer with lysozyme (a known bactericidal protein). CFU counts and OD_600_ were measured every two hours (Figure 8).

**Figure 8.**
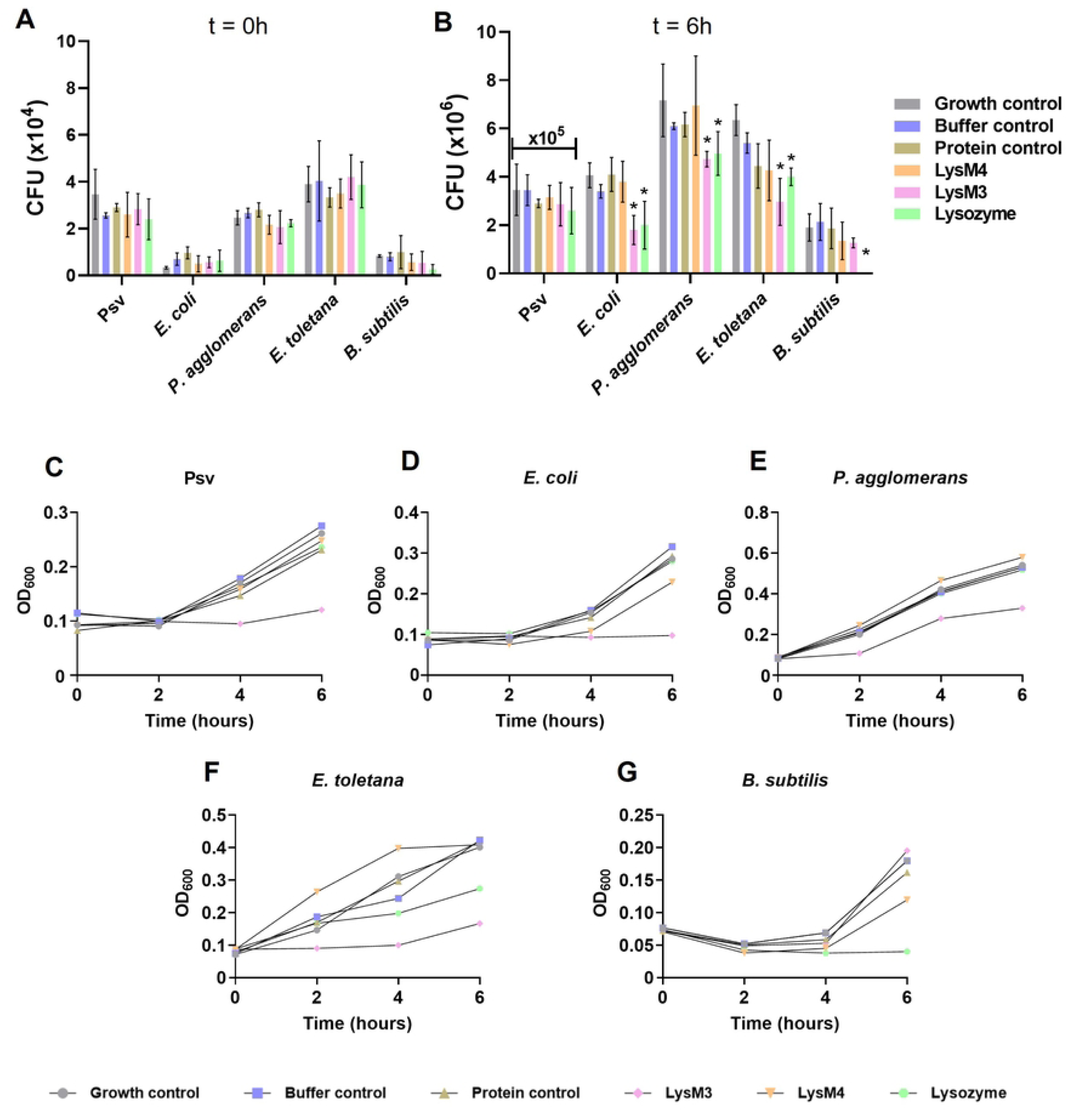
Bacteriostatic activity of LysM3 against Gram-negative bacteria. (A) Colony-forming units (CFU) recovered at time 0 from the indicated bacterial strains under the specified treatments. All cultures were adjusted to an initial optical density (OD₆₀₀) of 0.05. The following treatments were tested for bacteria grown in KB medium: Growth control, no additions; buffer control, supplemented with buffer only; protein control, as a control protein with no effect on bacterial growth, a transcriptional regulator from the AraC family was used; LysM4, purified LysM4 (contains only LysM domain); LysM3, purified LysM3; lysozyme, from chicken egg white. Data normality was assessed using the Shapiro-Wilk and Kolmogorov-Smirnov tests, and statistical significance was determined using Student’s t-test (p < 0.05). (B) CFU recovered after 6 hours of incubation at 28°C under the same treatments. Significant differences between treatments (p < 0.05) are indicated by an asterisk (*). (C-G) Growth curves showing OD₆₀₀ over 6 hours of incubation for each bacterial strain (indicated in each panel) under the treatments indicated below.

At the start of the assay (Figure 8A), CFU counts showed no statistically significant differences among treatments for any strain, confirming similar initial conditions. After 6 hours of incubation (Figure 8B), significant reductions in CFU counts were observed in the suspensions of all Gram-negative strains (*E. coli, P. agglomerans,* and *E. toletana*) treated with LysM3 or lysozyme, compared to controls. In contrast, no significant differences were detected for Psv NCPPB 3335 or *B. subtilis*, except for the lysozyme treatment, which completely inhibited *B. subtilis* growth.

Growth curves corroborated the trends observed in CFU counts. For Psv NCPPB 3335 and *E. coli* DH5α (Figures 8C, D), cultures treated with LysM3 exhibited slower growth compared to controls. Similarly, LysM3 significantly delayed growth progression for *P. agglomerans* DAPP-PG 734 and *E. toletana* DAPP-PG 735 (Figures 8E, F). In contrast, only the treatment with lysozyme caused a significant decrease in OD_600_ for *B. subtilis* 3610 (Figure 8G). These findings suggest that LysM3 exhibits bacteriostatic activity specifically against Gram-negative bacteria, suggesting its potential as an antimicrobial agent.

## Discussion

In this study, we characterized the secretome of *P. savastanoi* pv. *savastanoi* NCPPB 3335 and identified five proteins containing LysM domains, which we show play critical roles in the physiology and virulence of this phytopathogenic bacterium. Three of these proteins (LysM1, LysM2, and LysM4) are putatively involved in T4P formation, while LysM3 and LysM5 likely function as PG hydrolases with a role in the homeostasis of the bacterial cell wall. The loss of function of LysM3 and LysM5 resulted in severe defects in cellular septation, reducing the ability of Psv to colonize plant tissues and cause disease symptoms. Additionally, we demonstrated that LysM3 exhibits inhibitory activity against Gram-negative bacteria such as *Pantoea agglomerans* and *Erwinia toletana*, but not against Gram-positive bacteria such as *Bacillus*. These findings highlight the importance of PG hydrolases in Psv pathogenicity and their potential role in bacterial competition within the tumor niche.

LysM1, LysM2, and LysM4 are involved in type IV pili assembly, which is likely to influence the ability of the bacterium to adhere to and colonize plant tissue (46). The loss of function of these proteins results in reduced tumor volume and diminished tissue colonization capacity, with the defective Δ*lysM1* strain showing the most pronounced effects. This is further supported by the significant reduction in swimming and twitching motility observed in the Δ*lysM1* mutant. Similarly, in *Pseudomonas syringae* pv. tabaci 6605, which infects herbaceous plants, mutants in type IV pili have been shown to exhibit a significant reduction in swimming motility in semisolid medium as well as decreased virulence (55). No differences were observed in competition assays, suggesting that the wild-type strain may compensate for the reduced motility of the mutants during the infection process. LysM1 is orthologous to proteins widely distributed in bacteria and classified within the FimV/HubP family polar landmark proteins, including FimV from *Pseudomonas aeruginosa* (WP_003163304.1) (42), HubP from *Shewanella putrefaciens* (ABP76163.1; WP_011919521.1) (56) and TapV from *Ralstonia pseudosolanacearum* (WP_011001922.1) (57). Although proteins of this family have different roles in different species, mutants lacking FimV, HubP and TapV are impaired in swimming and/or twitching motility as well as, for FimV and TapV, severely reduced virulence. These phenotypes are mirrored by a LysM1 mutant, supporting the conservation of functions across taxa. Loss of type IV pili or of flagellar motility was shown to cause a reduction in virulence in a *P. syringae* pathovar phylogenetically very close to Psv, mainly because of alterations in the regulation of the expression of virulence-related genes (58). It is therefore likely that the activity of LysM1 is impacting the expression of diverse virulence genes, which should be further investigated. Importantly, mutants in LysM1 and in its paralog LysM2 (41.5% global similarity) displayed different phenotypes, as it occurs with the corresponding homologs in *R. pseudosolanacearum* (57). These phenotypic differences may be due to the presence of the tetratricopeptide repeat (TPR) domain, found exclusively in the *lysM1* product. This domain mediates protein-protein interactions (41) and is often associated with bacterial motility (56,59,60). Twitching motility, which depends on type IV pili, requires pili assembly, extension, and retraction. TPR domains may coordinate these interactions and potentially regulate flagellar-associated proteins, though their role in swimming motility is less clear (59,60). This could explain the reduced twitching and swimming motility in the LysM1 mutant.

In turn, LysM4 is an ortholog of protein TsaP, which was shown to be important for the correct assembly of the type IV pili by anchoring the secretin complex to the peptidoglycan layer (61,62). Mutants in TsaP did not show growth defects in *Mixococcus xanthus* (61) or in twitching motility in *P. aeruginosa* (62), as it occurs with Psv mutants in *lysM4*, but its overexpression in *P. aeruginosa* induced the cyclic di-GMP signal cascade, suggesting a role in regulation. Future studies should include adhesion assays to better understand the role of these proteins in tissue colonization. Additionally, generating double mutants could help clarify potential functional redundancies between these LysM proteins and reveal more pronounced phenotypes related to pili assembly, motility, or tissue colonization.

LysM3 and LysM5 are orthologs of proteins NlpD and AmiC, which participate in septal PG splitting during cell division (63,64). Our results indicate that LysM3 and LysM5 are also essential for cell division in Psv, as the Δ*lysM3* and Δ*lysM5* mutants display a filamentous phenotype characterized by cells remaining connected at their polar ends. This phenotype is consistent with defects in cellular septation, suggesting the conservation of function for these proteins. Previous research showed that NlpD did not have PG-hydrolase activity in *E. coli* and *Neisseria gonorrhoeae*, but rather acted as a potent and specific activator of the N-acetylmuramyl-l-alanine amidase AmiC, one of the three PG hydrolases playing a vital role in cell separation (63,65). However, the differential phenotypes shown by the *ΔlysM3* and *ΔlysM5* mutants (Figures 5 and 6), indicate that LysM3/NlpD is carrying out additional functions in Psv besides its predicted role activating LysM5/AmiC. This is not surprising, because this protein has been found to be involved in different phenomena in diverse bacterial species (66–69). These findings suggest that LysM3 and LysM5 could be promising targets for the development of antimicrobial strategies aimed at bacterial cell division.

The loss of function of LysM3 and LysM5 not only affected cell division but also significantly reduced the ability of Psv to colonize plant tissues and cause disease symptoms. The Δ*lysM3* and Δ*lysM5* mutants showed reduced tumor formation in olive plants and decreased competitiveness against the wild-type strain. These results align with studies in other plant or animal bacterial pathogens, such as the *Pseudomonas syringae* group (to which Psv belongs), *Ralstonia solanacearum* and *Yersinia pestis* where NlpD is implicated in virulence, host niche adaptation, biofilm formation, motility, iron acquisition, and/or activity of the twin-arginine system (66,69,70).

Previous research predicted that NlpD of *E. coli* did not bind zinc, due to its degenerated M23 domain that, as it occurs with LysM3 (Supplementary Figure 3), only possesses two of the four active-site residues in the HxxxD and HxH motifs (63). However, nanoDSF assays strongly suggest that LysM3 does interact with Zn^2+^, as incubation with the metal caused an increase of nearly 20°C in the protein stability (Figure 7). These assays also indicated the interaction of LysM3 with peptidoglycan fragments, such as D-Ala-D-Ala and mDAP, suggesting that this protein may recognize peptide moieties of the PG layer. Purified NlpD from *Neisseria gonorrhoeae* was shown to also bind purified PG, although, as it occurs with NlpD from *E. coli*, did not degrade PG in solution (63,65). Additionally, the interaction of LysM3 with oligosaccharides such as NAG_4_ and NAG_6_ suggests that it may also recognize chitin fragments, expanding its potential role in host interactions. Studies in *Agrobacterium tumefaciens* have shown that PG hydrolases release peptidoglycan fragments that are recognized by plant immune receptors, triggering defense responses (9). Future studies should investigate whether LysM3 modulates host immune responses during infection.

LysM3 exhibits an intriguing inhibitory activity exclusively against Gram-negative bacteria, such as *Pantoea* and *Erwinia*, suggesting a potential role in bacterial competition within the tumor niche. This phenomenon is reminiscent of other bacteria, like *Pseudomonas aeruginosa*, where peptidoglycan hydrolases act as bactericidal weapons to eliminate competitors (71). The specificity of LysM3 aligns with the described variability in the targets of M23 domain-containing proteins and their highly specific capacity to hydrolyze peptidoglycan (43). NanoDSF assays support the hypothesis that LysM3 is specific to the peptidoglycan of Gram-negative bacteria, as the greatest destabilization was observed in the presence of mDAP, a distinctive component of their peptidoglycan (43).

Interestingly, despite the more complex cell envelope of Gram-negative bacteria, as compared to Gram-positive organisms, LysM3 appears to only interact with the Gram-negative bacterial cell wall. This interaction may occur during the cell division phase of *P. agglomerans* and *E. toletana*, when the peptidoglycan layer could be temporarily weakened or remodeled, making it more accessible (72). This suggests a potential mechanism by which LysM3 could overcome the structural barriers of Gram-negative bacteria, exerting its inhibitory effect at a critical moment. Furthermore, such activity might provide a competitive advantage to *Pseudomonas savastanoi* during infection. Further studies are required to decipher the role of this potential inhibitory activity in bacterial competition and its implications for the dynamics within the tumor niche.

Although this work provides new insights into the role of PG hydrolases in Psv physiology and virulence, several questions remain. For instance, it is unclear how the expression of LysM3 and LysM5 is regulated during infection or whether these proteins have specific roles at different stages of the bacterial life cycle. Additionally, the molecular mechanisms underlying the inhibitory activity of LysM3 against Gram-negative bacteria requires further characterization. Future studies could explore whether this activity is mediated by peptidoglycan hydrolysis or by the release of fragments that compromise cell-wall integrity. Finally, the potential biotechnological application of LysM3 as a biocontrol agent against bacterial pathogens in crops warrants further investigation.

In summary, this study demonstrates that proteins LysM3 and LysM5 are essential for cell division, fitness, and virulence in *P. savastanoi*. Moreover, the inhibitory activity of LysM3 against Gram-negative bacteria suggests a role in bacterial competition within the tumor niche. These findings not only enhance our understanding of pathogenicity mechanisms in phytopathogenic bacteria but also open new avenues for developing disease control strategies in crops.

## Material and Methods

### Bacterial strains, media, and growth conditions

The bacterial strains, plasmids, and primers used in this study are listed in Supplementary Tables S2 and S3. *Pseudomonas savastanoi* and *Escherichia coli* were routinely grown at 28°C and 37°C, respectively, in lysogeny broth (LB) medium (73) without glucose and containing 0.5% NaCl. For induction experiments, *P. savastanoi* was maintained in HIM medium for 12 or 24 h (39) at 28°C. Growth inhibition and motility assays were carried out in Kinǵs B medium (KB) (74). When required, antibiotics were added to the media at the following final concentrations: for *P. savastanoi,* ampicillin (400 µg/mL), gentamicin (10 µg/mL), kanamycin (7 µg/mL), nitrofurantoin (25 µg/mL), and cycloheximide (100 µg/mL); for *E. coli*, ampicillin (100 µg/mL), gentamicin (10 µg/mL) and kanamycin (50 µg/mL).

### Isolation of secretomes

The secretome of the wild-type strain Psv NCPPB 3335 and its mutants Psv-ΔhrpA and Psv-ΔhrpL were isolated as follows. First, strains were grown overnight in LB medium at 28°C with shaking. The following day, cultures were diluted to an OD_600_ of 0.05 in fresh LB medium and incubated at 28°C with shaking until reaching an OD_600_ of 0.5. The cultures were then centrifuged, and the cell pellets were washed three times with Hrp-inducing medium (HIM)(39), resuspended in the same volume of medium HIM and incubated at 28°C with shaking for 12 or 24 hours. Following incubation, the cultures were centrifuged, and the supernatants were collected and filtered through 0.22 μm filters. The filtered supernatants were concentrated using Amicon^®^ Ultra filters (MercK Millipore Ltd., Ireland) and subjected to gel-assisted proteolysis for peptide extraction.

### Gel-assisted digestion and peptide extraction of culture supernatants

Gel-assisted proteolysis was performed by trapping the protein solutions in a polyacrylamide gel matrix. Briefly, 45 μL of each sample were mixed with 14 μL of 40% acrylamide monomer solution, 2.5 μL of 10% ammonium persulfate, and 1 μL of *N,N,N′,N′*-tetramethylethylenediamine, and the mixture was allowed to polymerize completely for 20 minutes at room temperature. After polymerization, the gel was cut into 1-2 mm cubes using a scalpel and treated with 50% acetonitrile (ACN) in 25 mM ammonium bicarbonate. The gel pieces were then dehydrated and dried with ACN.

For protein reduction, the gel pieces were incubated with 10 mM dithiothreitol (DTT) in 50 mM ammonium bicarbonate for 30 minutes at 56°C. Excess DTT was removed, and cysteine residues were carbamidomethylated by incubating the gel pieces with 55 mM iodoacetamide in 50 mM ammonium bicarbonate for 20 minutes at room temperature in the dark. After carbamidomethylation, the gel pieces were dehydrated again.

Protein digestion was performed by rehydrating the gel pieces in 10 ng/μL trypsin solution (Promega, USA) and incubating them at 30°C overnight. Peptides were extracted from the gel pieces with 0.1% formic acid (FA) in ACN for 30 minutes at room temperature. The samples were then dried in a SpeedVac^TM^ vacuum concentrator to remove residual ACN and ammonium bicarbonate. The dried peptides were reconstituted in 0.1% FA, treated with ultrasound for 3 minutes, and centrifuged at 13,000 x *g* for 5 minutes. Finally, the samples were purified and concentrated using C18 ZipTip^®^ pipette tips (MercK Millipore Ltd., Ireland) according to the manufacturer’s instructions and transferred to injection vials for analysis.

### Liquid chromatography and mass spectrometry

The samples were analyzed using an Easy nLC 1200 UHPLC system coupled to a hybrid quadrupole-linear trap-Orbitrap Q-Exactive HF-X mass spectrometer (Thermo Fisher Scientific, USA). Data acquisition and instrument operation were performed using Tune 2.9 and Xcalibur 4.1.31.9 software. The mobile phases for UHPLC consisted of solvent A (0,1% FA in water) and solvent B (0.1% FA in 80% ACN).

From a thermostatized autosampler, 1 µL (100 ng) of the peptide mixture was loaded onto a precolumn (Acclaim PepMap 100, 75 µm x 2 cm, C18, 3 µm, 100 Å, Thermo Fisher Scientific, USA) at a flow rate of 20 µL/min. Peptides were then eluted through an analytical column (PepMap RSLC C18, 2 um, 100 Å, 75 um x 25 cm, Thermo Fisher Scientific, USA) using a 120-minute gradient from 5% to 20% solvent B, followed by a 5-minute gradient from 20% to 32% solvent B, and finally a 10 minutes gradient to 95% solvent B. The column was re-equilibrated with 5% solvent B at a constant flow rate of 300 nL/minute. Prior to sample analysis, external instrument calibration was performed using the LTQ Velos ESI Positive Ion Calibration Solution (Pierce, IL, USA). Internal calibration was achieved using the polydimethylsiloxane ion signal at m/z 445.120024 from ambient air.

For mass spectrometry analysis, MS1 scans were acquired in the m/z range of 375-1600 at a resolution of 120,000. In data-dependent acquisition mode, the 15 most intense precursor ions with charges of +2 to +5 were isolated within a 1.2 m/z window and fragmented to obtain corresponding MS2 spectra. Fragmentation was performed in a high-energy collisional dissociation (HCD) cell with a fixed first mass at 110 m/z, and the resulting ions were detected in the Orbitrap mass analyzer at a resolution of 30,000. Dynamic exclusion was set to 30 seconds. The maximum ion accumulation times were 50 ms for MS and 70 ms for MS2 scans. Automatic gain control (AGC) was used to prevent overfilling of the ion trap, with targets settled to 3 × 10^6^ ions for full MS scans and MS2 scans and 2 × 10^5^ ions for MS2 scans.

### Protein identification

MS/MS spectra were searched against the *P. savastanoi* pv. savastanoi database (UniProt proteome ID: UP000005729; 4934 sequences). Raw data were analyzed using the Proteome Discoverer 2.5 software (Thermo Fisher Scientific, USA) with the Sequest HT search engine. Precursor and fragment ion mass tolerances were set to 10 ppm and 0.02 Da, respectively. Two missed trypsin cleavage sites were allowed.

Variable modifications included a methionine oxidation and N-terminal acetylation, while carbamidomethylation of cysteine residues was set as a fixed modification. The false discovery rate (FDR) for peptide and protein identification was determined using the Percolator software (integrated into Proteome Discoverer 2.5) based on a target-decoy approach, with a decoy database generated by inverting the protein sequences. A strict FDR threshold of 1% was applied. Only proteins identified by at least two unique peptide sequences were accepted.

### Screening and classification of proteins found in the secretome

The bioinformatic workflow SecretFlow, designed for the selection of secreted proteins, integrates multiple software tools. Proteins containing a signal peptide were identified using SignalP 5.0 (75), PrediSi (76), and TATFind 1.4 (77). Proteins with hydrophobic regions were filtered out using Phobius (78) and TMHMM 2.0 (79). The subcellular localization of the selected proteins was predicted using the SOSUI-GramN database (80). Venn diagrams were generated to visualize overlapping protein sets using the Good Calculators website (https://goodcalculators.com/venn-diagram-maker/).

Ortholog identification was performed using sequence searches in eggNOG 6.0.0 (81), and verified for individual bacterial strains through reciprocal blastp analysis on the NCBI server (https://blast.ncbi.nlm.nih.gov/Blast.cgi) against their complete proteomes.

### Functional enrichment analysis

Functional enrichment analysis was performed using the ShinyGO web platform (version 0.80; http://bioinformatics.sdstate.edu/go/) to identify the roles of genes of interest identified in secretome analyses. The list of genes was formatted with compatible identifiers and uploaded to the ShinyGO interface. The *P. savastanoi* pv. phaseolicola 1448A database was selected to ensure accurate genomic annotation matching.

Enrichment analysis was conducted for the following categories: Gene Ontology Molecular Function (MF), Gene Ontology Biological Process (BP), Uniprot, and InterPro database. Statistical significance was assessed using the Benjamini-Hochberg method to control the FDR at a threshold of 0.05. Comparative functional enrichment graphs were generated using Python.

### Prediction of functional gene relationships

Functional relationships between gene products were predicted using the STRING database (https://string-db.org/). A custom database was created using the proteome of Psv NCPPB 3335. Each LysM protein was analyzed individually to identify potential interactions and functional associations.

### Construction of *P. savastanoi* mutants and complemented strains

The *lysM* genes were deleted from Psv NCPPB 3335 using the plysMX-Km plasmid collection, which contain DNA fragments flanking each *lysM* gene (approximately 1.2 kb on each side) separated by an *nptII* gene conferring kanamycin resistance. This procedure followed previously established methods (82).

For the construction of complemented strains, the complete coding sequence of each *lysM* gene, including its predicted ribosome-binding site (RBS), was amplified from Psv NCPPB 3335 by PCR. The resulting amplicons were verified by sequencing and individually cloned into plasmid pBBR1MCS-5 (Gm^R^) under the control of the *P_lac_*promoter.

### Plant Assays

Pathogenicity assays were conducted on *Olea europaea* plants grown *in vitro* and *ex vitro*. Plants were derived from seeds of the cv. Arbequina germinated *in vitro* and maintained at 60% humidity and 26°C.

For *in vitro* virulence assays, a single wound was made in the stem of each plant, and 2 µL of the bacterial suspension (1 × 10^7^ CFU/mL) was inoculated into the wound. For *ex vitro* virulence assays, two wounds were made in the stem of each plant using a scalpel, and each wound was inoculated with 20 µL of the bacterial suspension (1 × 10^7^ CFU/mL). Four plants were used per strain in each of the three replicate assays.

In the *in vitro* assay, tumor volume was measured at the end of the 30-day trial using a 3D scanner and the Blender 3.5 software (Blender Institute, Amsterdam, The Netherlands). In the *ex vitro* assay, tumor volume was measured weekly over a 57-day period as described (83,84). At the end of the trial, symptoms were photographed using a high-resolution digital camera (Nikon DXM 1200). Bacteria were recovered from tumors by grinding the tissue in 1 mL of sterile 10 mM MgCl_2_ using a mortar and pestle. Serial dilutions were plated onto LB medium supplemented with nitrofurantoin, cycloheximide, and the appropriate antibiotic, as previously described (34,37,85).

Competitive index (CI) assays were performed on *in vitro* olive plants. Plants were inoculated with a 1:1 mixture of wild-type and mutant bacterial suspensions prepared in 10 mM MgCl_2_. A single wound per plant was inoculated with approximately 5 × 10³ CFU of each strain, using five plants per experiment. After 30 dpi under controlled conditions (25°C, 50-60% humidity, and 16-hour light photoperiod), symptoms were visualized using a stereo microscope (Leica MZ FLIII; Leica Microsystems, Wetzlar, Germany). Bacteria were recovered from tumors as described above, and serial dilutions were plated onto LB medium, and LB supplemented with kanamycin (LB+Km). Colony counting was performed after 2 days of incubation at 28°C. The number of CFU corresponding to the mutant strain was determined from colonies grown on LB+Km plates, while the number of wild-type bacteria was calculated by subtracting the mutant CFU from the total CFU grown on LB plates (86,87). The CI was calculated as the ratio of mutant to wild-type bacteria in the output sample divided by the ratio of mutants to wild-type bacteria in the input (inoculum) sample (88,89).

### Motility assays

Twitching and swimming motility assays were conducted using KB medium containing 0.3% (wt/vol) and 1% (wt/vol) agar, respectively, at 25°C. Twitching motility was evaluated following previously established protocols (57,90) with minor modifications. Briefly, bacterial biomass was resuspended in 10 mM MgCl_2_, and 2 µl of the suspension (OD_600_ of 0.1) was deposited onto 1 ml KB agar overlaid on slides. Coverslips were placed over the inoculated medium to allow twitching motility at the interstitial interface created, and the samples were incubated for 24 h in a Petri dish under humid conditions. After incubation, the twitching areas were examined using a Nikon Eclipse E800 light microscope with a 20x objective lens. For swimming assays, bacterial biomass was resuspended in 10 mM MgCl_2_, washed three times, and adjusted to an OD_600_ of 1. Then, 3 µl of the bacterial suspension was inoculated into the center of the KB agar medium. After 3 days of incubation, swimming halos were measured, and their area was calculated. Statistical analyses were performed using GraphPad Prism 8 software. Images were captured using a Panasonic Lumix camera.

### Growth curve analysis

Bacterial growth curves were determined by inoculating *P. savastanoi* cultures in 50 mL of LB broth in 250 mL flasks. Cultures were incubated at 28°C with shaking at 150 rpm. Each culture started with an initial OD_600_ of 0.05, and growth was monitored by measuring OD_600_ at hourly intervals using a spectrophotometer. To complement optical density measurements, serial dilutions were prepared in 10 mM MgCl_2_ and plated onto LB agar every hour to enumerate CFU. The experiment was performed in triplicate to ensure reproducibility. Statistical analysis was conducted using GraphPad 8 Prism software.

### Confocal microscopy

Confocal microscopy observations were conducted using a Zeiss LSM 880 confocal microscope. A 100 µL volume from an overnight culture of each mutant was prepared for imaging. Cells were stained with FM4-64, a membrane dye, at the manufacturer-recommended concentration, and incubated for 5 minutes at room temperature. Images were captured and processed using the ZEN Blue software.

### Flow cytometry

Flow cytometry measurements of size and complexity were conducted on cultures in the exponential growth phase in LB medium for all analyzed strains using a FACSverse flow cytometer. Up to a maximum of 10,000 events were counted in 1 mL of undiluted culture. The experiment was repeated three times, and data was processed and analyzed using the Kaluza software.

### Protein induction and purification

The expression vectors (pET28a::lysM3 and pET28a::lysM4) were transformed in calcium chloride-treated *E. coli* BL21(DE3) cells. Overexpression was induced by adding 200 mM isopropyl-β-D-thiogalactoside (IPTG) when the culture reached an OD_600_ of 0.5-0.8 and continuing incubation at 18°C for 24 h in LB medium containing 50 µg/mL kanamycin. Cells were harvested by centrifugation at 4,500 rpm for 30 min at 4°C. For purification, cell pellets were resuspended in lysis buffer (50 mM Tris-HCl pH 8.0, 500 mM NaCl, 10 mM imidazole, 10 mM DNAse, 1 mM MnCl_2_, 10 mM MgCl_2_, 4 mM β-mercaptoethanol) and incubated at room temperature for 15 min. The cell suspension was lysed by sonication for 15 min (15 s pulses at 40% power) under ice cooling using an Ultrasonic processor (Sigma-Aldrich). The crude lysate was cleared by centrifugation at 14,000 rpm for 15 min at 4°C and purified by immobilized metal affinity chromatography (IMAC) using a HisTrap HP 1 mL column (Cytiva Europe GmbH, Freiburg in Breisgau, Germany) on an ӒKTA Go FPLC device (Cytiva). After washing with five column volumes (CV) of IMAC buffer A (50 mM Tris-HCl pH 8.0, 500 mM NaCl, 10 mM imidazole, 4 mM β-mercaptoethanol), LysM proteins were eluted with a linear gradient of IMAC buffer B (50 mM Tris-HCl pH 8.0, 500 mM NaCl, 500 mM imidazole, 4 mM β-mercaptoethanol). Eluted proteins were concentrated using Amicon^®^ Ultra filters (MercK Millipore Ltd., Ireland) and subjected to a size-exclusion chromatography using a HiLoad 16/600 Superdex 200 pg (Cytiva) column equilibrated in SEC buffer (20 mM Tris-HCl pH 8.0, 150 mM NaCl, 1 mM DTT), fractions containing LysM proteins were concentrated, flash-frozen in liquid nitrogen and stored at −80°C.

### NanoDSF assays

Thermoshift assays were carried out to estimate the inflection temperature (*T*_i_) associated with unfolding transitions of LysM3 in the presence and in the absence of ligands. Experiments were performed using the Tycho NT.6 nanoDSF instrument (NanoTemper Technologies GmbH, München, Germany) with manufacturer-designed capillaries. DSF experiments were set up in a final volume of 10 µL in DSF buffer (20 mM Hepes pH 8.0, 150 mM NaCl) containing 5 µM of protein and, when required, 0.5 mM ZnCl_2_ and the following ligands: D-alanine-D-alanine at 5 mM, mesodiaminopimelic acid at 5 mM, diacetyl-L-lysine-D-alanine-D-alanine at 1 mM, N-acetyl-muramoyl-L-alanine-D-isoglutamine at 20 mM, tetra-N-acetylglucosamine at 1 mM and hexa-N-acetylglucosamine at 1 mM. Three independent runs were used in each case to calculate mean *T*_i_ values and the corresponding standard deviations. The data was plotted using GraphPad Prism 8.

### Protein structure analysis

Structural analyses were performed using the crystallographic structures of P60 from *Thermus thermophilus* (P60 Hexanag PDB code: 4zu3)(49), Csd1 from *Helicobacter pylori* (Csd1 PDB code: 5j1l)(50) and Pgp3 from *Campylobacter jejuni* (Pgp3:mDAP-D-Ala PDB code: 6jn1)(51) along with a model of LysM3 predicted by AlphaFold (91). Structural comparisons and alignments were performed using the Research Collaboratory for Structural Bioinformatics (RCSB) PDB pairwise structure alignment tool (92). Protein sequence alignment and structure visualization were carried out using Jalview (93) and PyMOL (Schrödinger, New York, United States), respectively.

### Growth inhibition assay

The bacterial strains used in this assay are listed in Supplementary Table 1. Overnight precultures were grown at 28°C with constant shaking. The following day, cultures were adjusted to a final OD_600_ of 0.05 and distributed in 270 µL aliquots by triplicate into a 96-well plate. Bacteria grown in KB medium were subjected to the following treatments by adding 30 µL of the corresponding solution, with final concentrations indicated in parentheses: growth control (no additions); buffer control (20 mM Tris pH 8, and 150 mM NaCl); protein control, consisting of a transcriptional regulator from the AraC family encoded in a Psv NCPPB 3335 genomic island (94), which has no effect on bacterial growth and was purified under the same conditions as LysM3 and LysM4 (15 µM); LysM4, purified LysM4 protein (15 µM); LysM3, purified LysM3 protein (15 µM); and lysozyme, lysozyme from chicken egg white (Apollo Scientific, Whitefield, United Kingdom) (15 µM). The plates were incubated at 28°C, and OD_600_ measurements were recorded at regular intervals. Additionally, samples were collected for serial dilutions in 10 mM MgCl_2_, plated onto KB agar plates, and incubated at 28 °C for two days. Data for OD_600_ and CFU counts were processed and analyzed using GraphPad Prism 8.

## Acknowledgements

We are grateful to A.I. Berrocal-Calle for her excellent technical assistance. We thank the *Servicios Centrales de Apoyo a la Investigación* (SCAI) of University of Málaga for the technical support. We are grateful to A. Arroyo-Mateo for kindly providing the protein control used in the growth inhibition assay. We are thankful to J.A. Hermoso for kindly providing the ligands used in the nanoDSF experiments. We also thank Bart Thomma for critical reading of the manuscript.

